# acorn: an R package for *de novo* variant analysis

**DOI:** 10.1101/2023.04.11.536422

**Authors:** Tychele N. Turner

**Author notes:** Correspondence to, Tychele N. Turner, Ph.D., Washington University School of Medicine, Department of Genetics, 4523 Clayton Avenue, Campus Box 8232, St. Louis, MO 63110.

## Abstract

**Background:** The study of *de novo* variation is important for assessing biological characteristics of new variation and for studies related to human phenotypes. Software programs exist to call *de novo* variants and programs also exist to test the burden of these variants in genomic regions; however, I am unaware of a program that fits in between these two aspects of *de novo* variant assessment. This intermediate space is important for assessing the quality of *de novo* variants and to understand the characteristics of the callsets. For this reason, I developed the R package acorn.

**Results:** acorn is an R package that examines various features of *de novo* variants including subsetting the data by individual(s), variant type, or genomic region; calculating features including variant change counts, variant lengths, and presence/absence at CpG sites; and characteristics of parental age in relation to *de novo* variant counts.

**Conclusions:** acorn is an R package that fills a critical gap in assessing *de novo* variants and will be of benefit to many investigators studying *de novo* variation.

## BACKGROUND

The study of *de novo* variants (DNVs) consists of 1) calling of DNVs from parent-child sequencing data, 2) quality control and DNV dataset characterization, and 3) assessment of DNVs for statistical interpretation (e.g., assessment of burden of DNVs in genomic regions in human phenotypes). Several tools exist for step 1 [1-4] and step 3 [5-9]. However, I am unaware of a tool that sits in between these steps at step 2. The analysis at step 2 is critical for inputting high quality data into step 3. I developed acorn as a strategy to centralize code I use for performing these quality checks for public distribution to help others also analyzing DNVs. The implementation of acorn as an R package is key for its use on multiple platforms. Functions are built so investigators can use them in several ways to assess their DNV data.

## IMPLEMENTATION

Acorn (version 0.99.5) is an R package with minimal prerequisites and is publicly available at https://github.com/TNTurnerLab/acorn. It requires baseline R packages graphics, stats, and utils making it a quick package to install by the user. Both source code and a pre-built version of the package are available allowing users to download and build the package themselves or quickly install using `R CMD INSTALL acorn_0.99.5.tar.gz`, respectively.

## RESULTS & DISCUSSION

### Acorn Functions

Acorn consists of several functions to analyze DNVs. One function is `readDNV`, which reads in DNV data and turns it into an R object for use with other functions within acorn. The next set of functions are used to extract different subsets of DNV data. These include `extractIndividual`, `extractSNVs`, `extractINDELs`, `extractMNVs`, `extractAutosomes`, `extractX`, and `extractY`. The `extractIndividual` function can extract an individual or set of individuals from the dataset. The `extractSNVs`, `extractINDELs`, and `extractMNVs` functions are used to extract variant types including single-nucleotide variants (SNVs), small insertions/deletions (INDELs), or multi-nucleotide variants (MNVs) from the dataset, respectively. Finally, the `extractAutosomes`, `extractX`, and `extractY` functions are all used to extract DNVs by genomic regions. The `extractAutosomes` function extracts the autosomes (chromosomes 1 to 22) from the dataset. The `extractX`, and `extractY` functions extract DNVs on the X and Y chromosomes, respectively.

The next set of functions are used for examining summary characteristics of DNV data. These include the `calculateTiTvRatio, calculateDeletionInsertionRatio, calculateDeletionLengths`, `calculateInsertionLengths`, and the `calculateMNVLengths` functions. The `calculateTiTvRatio` function automatically grabs only the SNVs from the DNV object for the calculation of the transition/transversion (Ti/Tv) ratio. It returns the counts of transitions, the counts of transversions, the Ti/Tv ratio,

and a barplot of the different types of SNV changes observed in the DNV object. The `calculateDeletionInsertionRatio` automatically grabs only the INDELs from the DNV object for the calculation of the deletion/insertion ratio. It returns the counts of deletions, the counts of insertions, and the deletion/insertion ratio. The `calculateDeletionLengths`, `calculateInsertionLengths`, and `calculateMNVLengths` functions determine the lengths of the deletions, insertions, and MNVS, respectively. For each, it also returns a barplot of the lengths.

Another key feature of DNV data, is the percent at CpG sites. The `annotateCpG` function annotates and summarizes CpG information for the DNV dataset. It extracts single-nucleotide variants (SNVs) and assigns whether they are at a CpG site or not. This function also requires a pre-computed rda file for the CpG sites in the genome of interest. I have made this available for b38 of the human genome at: https://data.cyverse.org/dav-non/iplant/home/tycheleturner/genomic_annotations/cpg_b38.rda. The function returns a DNV object containing only SNVs and includes a column with a note on whether the DNV is at a CpG or not. This function also prints out the number of CpG and the percent of DNV SNVs at CpG.

The `countsPerIndividual` function returns the mean of the DNV counts per individual, the standard deviation of the DNV counts per individual, a plot of the density of the DNV counts per individual,and an object consisting of the sample name and the counts of their DNVs that can be assigned to another object.

The final set of functions focus on the parental age characteristics of DNVs. The first is `parentalAgeObject`, which takes in a counts object that is either the result of the `countsPerIndividual` function or is already read into an object from a file. This function returns back an object with the *de novo* counts and parental age data together. The fields in this file are sample, dnm_counts, fatherAge, and motherAge. The `parentalAge` function calculates the correlation between father’s and mother’s age at birth and DNV counts per individual, the results of the linear model taking the form: `lm(formula = dnm_counts ∼ fatherAge+motherAge, data = parentalAgeObject)`. The input required is output from the `parentalAgeObject` function in this package. This function returns the results of the linear model and a plot of father’s and mother’s age at birth and DNV counts. The `fatherAge` and `motherAge` functions are similar to the `parentalAge` function except that they only consider the father’s age or mother’s age, respectively.

### DNV Assessment with Acorn

We provide test data with acorn, on our acorn GitHub site, consisting of a subset of our DNV dataset from our recent paper [10] (dnms_from_Ng_et_al_2022_Human_Mutation_paper.txt.gz, Supplemental File 1), simulated count data (dnm_count_example.txt, Supplemental Table 1), and simulated parental age data (parental_age_example.txt, Supplemental Table 2). We detail the use of these in the Vignette included with the acorn package. Briefly, the DNV data consists of 9,741 DNVs of which 8,558 are SNVs and 1,183 are INDELs. For SNV changes (Figure 1A), the number of transitions is 5,708 and the number of transversions are 2,850 with a Ti/Tv ratio of 2.0. The number of deletions is 540 and the number of insertions is 643 with a deletion/insertion ratio of 0.8. Acorn also shows the length of deletions (Figure 1B) and insertions (Figure 1C). These are just a few ways that acorn can be used to examine DNV data.

**Figure 1:**
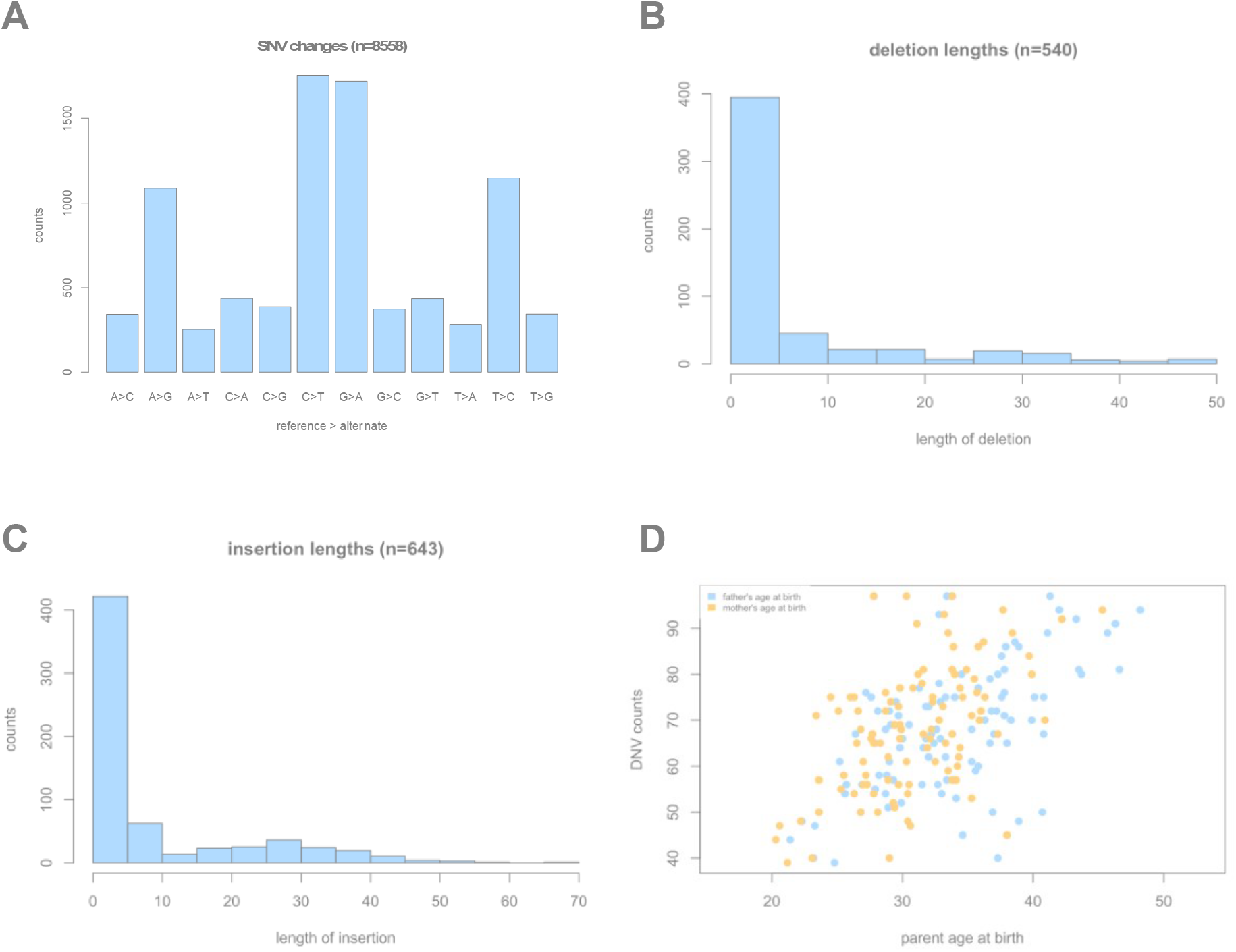
Example results from acorn. A) SNV changes as shown using the `calculateTiTvRatio` function, B) Deletion lengths from the `calculateDeletionLengths` function, C) Insertion lengths from the `calculateInsertionLengths` function, and D) DNV counts versus parental age at birth from the `parentalAge` function.

For the parental age analysis on the simulated data, acorn presents a plot of DNV counts versus parental age at birth (Figure 1D). It also shows the result of using a linear model to examine DNV counts versus the parental age at birth reporting back the coefficients, r-squared values, and confidence intervals. In the test set, the number of DNVs rises by 1.0 [0.5, 1.6] each increasing year of father’s age at birth and rises by 0.7 [0.1, 1.3] for each increasing year of mother’s age at birth.

## CONCLUSIONS

The summary statistics and metrics generated using acorn will help users to determine the quality of their DNV callset(s) and whether they should move forward with advanced statistical tests. Acorn fills a gap in genomic DNV analyses between the calling of DNVs and ultimate downstream statistical assessment (e.g., DNV association with phenotypes).

## Supporting information

dnms_from_Ng_et_al_2022_Human_Mutation_paper.txt.gz

dnm_count_example.txt

parental_age_example.txt

## AVAILABILITY AND REQUIREMENTS

**Project name:** acorn

**Project home page:** https://github.com/TNTurnerLab/acorn

**Operating system(s):** Platform independent

**Programming language:** R **Other requirements:** R 3.5.0 **License:** MIT

**Any restrictions to use by non-academics:** none

### LIST OF ABBREVIATIONS

DNV: *de novo* variant

## DECLARATIONS

Availability of data and materials: all data generated or analyzed during this study are included in this preprint in the supplementary information files and/or at the acorn GitHub site at https://github.com/TNTurnerLab/acorn, funding: this work was supported by grants from the National Institutes of Health (R01MH126933) and the Simons Foundation (Award #734069) to T.N.T., authors’ contributions: T.N.T. wrote the acorn R package, performed analyses, and wrote the paper.

## Notes

https://github.com/TNTurnerLab/acorn

### Competing Interest Statement

The authors have declared no competing interest.

